# Osteocyte death and bone overgrowth in mice lacking Fibroblast Growth Factor Receptors 1 and 2 in mature osteoblasts and osteocytes

**DOI:** 10.1101/474502

**Authors:** Jennifer McKenzie, Craig Smith, Kannan Karuppaiah, Joshua Langberg, Matthew J. Silva, David M. Ornitz

## Abstract

Fibroblast Growth Factor (FGF) signaling pathways have well established roles in skeletal development, with essential functions in both chondrogenesis and osteogenesis. In mice, previous conditional knockout studies suggested distinct roles for FGF receptor 1 (FGFR1) signaling at different stages of osteogenesis and a role for FGFR2 in osteoblast maturation. However, the potential for redundancy among FGFRs and the mechanisms and consequences of stage-specific osteoblast lineage regulation were not addressed. Here, we conditionally inactivate *Fgfr1* and *Fgfr2* in mature osteoblasts with an Osteocalcin-Cre or Dentin matrix protein 1-CreER driver. We find that young mice lacking both receptors or only FGFR1 are phenotypically normal. However, after 6 weeks of age these *Fgfr1/Fgfr2* double- and *Fgfr1* single-conditional knockout mice develop a high bone mass phenotype with increased periosteal apposition, increased endocortical woven bone with increased porosity, and biomechanical properties that reflect increased bone mass but impaired material properties. Histopathological and gene expression analyses show that this phenotype is preceded by a striking loss of osteocytes, and gradual activation of the Wnt/βCatenin signaling pathway. These data identify a role for FGFR1 signaling in mature osteoblasts/osteocytes that is required for osteocyte survival during postnatal bone growth.

## Introduction

Osteoprogenitors in the periosteum and bone marrow give rise to matrix producing osteoblasts. As bone grows, osteoblasts can become quiescent bone lining cells, die through apoptosis, or differentiate into osteocytes, long-lived cells that become embedded within the mineralized bone matrix.^(1,2)^ Although embedded within mineralized bone, osteocytes maintain communication with the periosteum, endosteum, and vasculature through a system of channels that form the osteocyte lacunocanalicular network.

Osteocytes function as sensory cells that respond to mechanical loading and other signals by releasing soluble factors that regulate osteoblast anabolic activity, mineral homeostasis, and osteoclast activity. Osteocytes express Sclerostin (*Sost*) and Dickkopf 1 (*Dkk1*) and osteoblasts express *Dkk2*, all potent inhibitors of Wnt signaling.^(3–10)^ In response to mechanical loading, expression of *Sost* and *Dkk1* is suppressed, at least in part through osteocyte production of prostaglandin E2.^(3–8,11,12)^ Interestingly, *Dkk1* and *Sost* reciprocally regulate each other’s expression; however, the anabolic activity of *Dkk1* is only revealed in the absence of *Sost*.^(13)^ Osteocytes also respond to parathyroid hormone (PTH), which likewise functions to suppress *Sost*, accounting for some of the anabolic effects of PTH.^(14)^ Osteocytes are a major source of the cytokine receptor activator of NF**k**B ligand (RANKL). Mechanical unloading and PTH signaling stimulate RANKL expression in osteocytes.^(14–17)^ Osteocytes are also a major source of FGF23, an endocrine member of the Fibroblast Growth Factor (FGF) family that primarily signals to the kidney proximal tubules to suppress phosphate resorption,^(18,19)^ but also signals to FGF receptor 3 (FGFR3) in osteocytes to inhibit mineralization.^(20)^

FGF signaling has well established roles in bone development with essential functions in both chondrogenesis and osteogenesis.^(21)^ In humans, mutations in FGF receptors (FGFRs) 1 and 2 that affect osteogenesis are found in Crouzon, Apert, Pfeifer, and Bent bone dysplasia syndromes. These mutations result in ligand dependent activation of the mutant FGFR and, in some cases, change in the ligand binding specificity of the FGFR. The only documented example of a FGFR loss of function mutation occurs in CATSHL syndrome, in which loss of FGFR3 results in skeletal overgrowth due to increased chondrogenesis.^(21,22)^

FGFRs are expressed in most, if not all, osteogenic lineages throughout development, and in mature bone. *Fgfr1* is expressed throughout limb bud mesenchyme at similar levels in condensing and surrounding cells. In condensing mesenchyme that will give rise to skeletal tissue, *Fgfr2* expression increases relative to the surrounding mesenchyme.^(23–28)^ Cells in the periphery of the mesenchymal condensation form the perichondrium and periosteum and express both *Fgfr1* and *Fgfr2*.^(24,27)^ In the definitive periosteum of developing bone, *Fgfr1* is expressed in mesenchymal progenitors and *Fgfr2* is expressed in differentiating osteoblasts.^(29–33)^ In the periosteum of mature bone, FGFR2 expression is lost, while FGFR1 expression persists in a subpopulation of perivascular cells, some of which express the pericyte marker NG2.^(34)^ Mature osteoblast and osteocytes also express FGFR3.^(20,35,36)^ During fracture repair, expression of both FGFR1 and FGFR2 are increased,^(37)^ and both FGFR1 and FGFR2 are expressed in periosteal derived mesenchymal stem cells.^(34)^ These expression patterns suggest the potential for both unique and redundant functions of *Fgfr1* and *Fgfr2* in the osteoblast lineage.

Conditional inactivation of *Fgf* ligands and receptors in skeletal lineages has been a successful approach to identify function during development and following injury. Conditional inactivation of *Fgfr1* and *Fgfr2* in limb bud mesenchyme (using *Prx1-Cre)*, or inactivation of *Fgfr1* in all mesenchyme (using *T(brachyury)-Cre*) causes severe skeletal hypoplasia,^(38,39)^ whereas inactivation of *Fgfr1* or *Fgfr2* in distal limb bud mesenchyme (using *AP2-Cre*) causes only a mild skeletal phenotype,^(40,41)^ suggesting redundancy between FGFRs 1 and 2 in distal limb mesenchyme. Conditional inactivation of *Fgfr1* in osteoprogenitor cells using *Col2a1-Cre* results in increased proliferation and delayed differentiation and matrix mineralization, and conditional inactivation of *Fgfr1* in committed osteoblasts using *Col1a1-Cre* causes enhanced mineralization and cortical bone thickening in 5- to 8-month-old mice.^(30)^ However, the mechanisms by which FGFR1 affects bone development and growth are not known. In contrast to *Fgfr1*, mice conditionally lacking *Fgfr2* in all bone lineages or harboring a mutation in the mesenchymal splice variant of *Fgfr2* (*Fgfr2c*), show impaired ossification, and decreased bone mineral density, suggesting that *Fgfr2* is a positive regulator of osteoblast maturation.^(24,42)^ The function of FGFR2 in mature osteoblast lineages is not known.

In this study, we investigated the bone-specific function of *Fgfr1* and *Fgfr2* in mature osteoblast/osteocyte-specific static and inducible conditional knockout mice. We find that mice lacking *Fgfr1* and *Fgfr2,* or *Fgfr1* alone, in mature osteoblasts develop a dramatic high bone mass phenotype that becomes histologically evident between 6 and 12 weeks of age. Bone pathology initiates as early as 3 weeks of age with the onset of osteocyte loss, which progresses to nearly complete loss by 12 weeks. This phenotype is recapitulated with a postnatal conditional knockout model that progresses with a similar time course. Biomechanical studies show that although whole-bone strength is increased in correlation with increased mass, bone material properties are diminished. These studies show that FGFR1 signaling in mature osteoblasts or osteocytes is essential for osteocyte viability and that the absence of FGFR1, either directly or secondary to osteocyte loss, increases bone formation.

## Materials and Methods

### Mice

Mice were group housed with littermates, in breeding pairs, or in a breeding harem (2 females to 1 male), with food and water provided ad libitum. Mice were housed in a pathogen-free facility and handled in accordance with standard use protocols, animal welfare regulations, and the NIH Guide for the Care and Use of Laboratory Animals. All studies performed were in accordance with the Institutional Animal Care and Use Committee at Washington University in St. Louis (protocol #20160113)

*Fgfr1^f/f^*,^(43)^ *Fgfr2^f/f^*,^(42)^ OC-Cre,^(44)^ Dmp1-CreER,^(45)^ *ROSA^Tomato^* (Ai9 allele),^(46)^ *ROSA^mTmG^*,^(47)^ mice have been previously described. Mice were of mixed sexes and maintained on a mixed C57BL/6J x 129X1/SvJ genetic background. Homozygous floxed alleles of *Fgfr1* and *Fgfr2* were maintained as double floxed mice (DFF) and outbred to hybrid C57BL/6J;129X1 mice every second generation and then backcrossed to homozygosity. OC-Cre; *Fgfr1^f/f^; Fgfr2^f/f^* mice are maintained by crossing OC-Cre; *Fgfr1^f/f^; Fgfr2^f/f^* males to *Fgfr1^f/f^; Fgfr2^f/f^* females, resulting in a 50% yield of experimental and *Fgfr1^f/f^; Fgfr2^f/f^*(DFF) mice (which serve as one type of control). A similar mating scheme was used to generate Dmp1-CreER; *Fgfr1^f/f^; Fgfr2^f/f^* mice. To induce Cre activation in Dmp1-CreER; *Fgfr1^f/f^; Fgfr2^f/f^* and *Fgfr1^f/f^; Fgfr2^f/f^* control mice, tamoxifen chow (400 mg/kg, Envigo Teklad, TD.130860) was given from 3 to 5 weeks of age.

### MicroCT and DXA Imaging

*In vivo* microCT was done on the right tibia and femur to evaluate cortical and cancellous bone measures over time (VivaCT 40, Scanco Medical, Wayne, PA; X-ray tube potential 70 kV, integration time 300 ms, X-ray intensity 114 µA, isotropic voxel size 21 µm, frame averaging 1, projections 1024, medium resolution scan). Measurements were made according to recommended standards,^(48)^ with cortical bone measured for 1 mm (48 slices) at a location 5 mm proximal to the tibia fibula junction: total area (T.Ar), bone area (B.Ar), cortical thickness (Ct.Th), tissue mineral density (TMD), and polar moment of inertia (pMOI). Metaphyseal bone was analyzed from a region 600 µm (28 slices) distal to the proximal growth plate in the tibia and proximal to the distal growth plate in the femur. Because of the blurred distinction between the cortical shell and the adjacent cancellous bone in the DCKO mice at ages > 12 weeks, we analyzed all tissue inside the periosteal margin (cortical shell + cancellous bone). Outcomes were: total volume (TV), bone volume (BV), bone volume to total volume ratio (BV/TV), and volumetric bone mineral density (vBMD). *In vivo* dual energy X-ray absorptiometry (UltraFocus100 DXA module, Faxitron, Tucson, Arizona) was used to measure percent fat, bone mineral content (BMC) and areal bone mineral density (aBMD) for the whole body of the mouse (excluding the head). To evaluate growth, body weight was recorded prior to each scan. Mice were scanned at 6, 12, 18, 24 and 32 weeks of age. For the Dmp1-CreER; *Fgfr1^f/f^; Fgfr2^f/f^* mice the bones were scanned *ex vivo* (X-ray tube potential 55 kV, integration time 300 ms, X-ray intensity 145 µA, isotropic voxel size 10 µm, frame averaging 1, projections 2048, high resolution scan), embedded in 2% agarose.

### Dynamic Histomorphometry

To mark bone formation, 12-week-old mice were injected with calcein green (10 mg/kg, IP, Sigma) and 5 days later with alizarin red (25 mg/kg, IP, Sigma), which was given 2 days prior to euthanasia. Tibia were embedded in poly-methyl methacrylate (plastic) and 100 μm thick cross-sections of the diaphysis were cut 5 mm proximal to the tibiofibular junction. Sections were mounted to glass slides and polished to a thickness of approximately 30 μm. To visualize the calcein and alizarin labels fluorescent images were taken (Olympus IX51 with Olympus DP70 camera or Leica TCS SPE Confocal). Following published standards^(49)^, the bone sections were analyzed for mineral apposition rate (MAR), mineralizing surface per bone surface (MS/BS) and bone formation rate per bone surface (BFR/BS) on the endocortical (Ec) and periosteal (Ps) surfaces (BIOQUANT OSTEO, BIOQUANT Image Analysis Corporation, Nashville, TN).

### Biomechanics

At 24 weeks age the mechanical properties of the femurs were evaluated with mechanical testing. Prior to testing, the midshaft of these bones (100 slices, 2.1 mm) were imaged with microCT to determine bone morphology (VivaCT 40, Scanco Medical, same parameters as above). After scanning, the femurs were prepared for three-point bending (span = 7 mm). Bones were preloaded to −0.3 N, then a displacement ramp of 0.1 mm/s was applied until failure (Dynamight 8841, Instron, Norwood, MA). As recommended by Jepsen et al.,^(50)^ the structural mechanical properties acquired from bending tests were stiffness, yield load, maximum load, post-yield displacement, and work-to-fracture. In combination with the bone morphology (moment of inertia) and beam theory, the estimated material properties of Young’s modulus, yield stress, and maximum stress were calculated. After whole-bone testing was completed, reference point indentation (RPI) testing was done at four locations near the midshaft (two on each side of the break). The parameters measured included first indentation distance (ID 1^st^), total indentation distance (TID), indentation distance increase (IDI), first unloading slope (US 1^st^), and average creep indentation distance (CID).

### Histology, immunohistochemistry, and immunofluorescence

For histological analysis of long bones, intact femurs and tibias were isolated, fixed in 4% PFA/PBS overnight at 4°C or fixed in 10% buffered formalin overnight at room temperature. Bones were rinsed in water several times and decalcified in 14% EDTA/PBS for 2 weeks. Paraffin-embedded tissue sections (5 μm) were stained with Hematoxylin and Eosin (H&E), tartrate resistant acid phosphatase (TRAP), von Kossa or Alizarin Red. The organization of lamellar and woven bone was qualitatively assessed with polarized light microscopy. Both brightfield and polarized light microscopic images were obtained at 20x using an Olympus BX51P Polarizing Microscope and a Leica EC3 Camera.

For immunohistochemistry, paraffin sections or cryosections were rehydrated and treated with 0.3% hydrogen peroxide in methanol for 15 min to suppress endogenous peroxidase activity. Antigen retrieval was achieved by microwaving the sections in 10 mM citrate buffer (pH 6.0) for 10 min followed by gradual cooling to room temperature. Sections were incubated overnight at 4°C with the following primary antibodies: DMP1 (Abcam, ab82351; 1:100), mSOST (RD systems, AF1589; 1:100). Secondary antibody was Alexa Fluor 488 donkey anti-rabbit (Life Technologies, A21206; 1:1000). Immunofluorescence imaging was performed on a Zeiss Apotome II fluorescence microscope.

TUNEL staining was performed using the DeadEnd Flurometric TUNEL system staining kit (Promega, G3250). All kit instructions were followed for paraffin-embedded tissues with a 5 min proteinase K treatment and fixation with 4% paraformaldehyde. Negative controls with rTdT Enzyme were run with each group. Imaging was performed on a Leica TCS SPE Confocal microscope.

### Determination of osteocyte density and viability

Osteocytes were histologically categorized as alive, dying, and dead based on their appearance on H&E stained cortical sections (see Figure 5J below). Alive cells were characterized by a whole nucleus with little to no empty space in the lacuna. Dying cells were identified morphologically as having either a fragmented nucleus or a small polarized nucleus within an otherwise empty lacuna. Dead cells were identified as having no hematoxylin-stained material in the lacuna. A small number of histological features (<10%) could not be assigned a definitive category and were not counted. These may represent histological sectioning artifacts or ossified lacunae. Osteocytes from all three categories were counted in the anterior tibial cortex in a 3 mm region beginning 1 mm from the growth plate. Osteocyte density was calculated by dividing the number of counted cells/lacunae by the area of cortical bone. Cortical bone area was quantified by measuring the total bone area in the analysis region and then subtracting the area of any region inside this total bone area that was not cortical bone, e.g. marrow space or other cavities.

### Real-time quantitative PCR (RT-qPCR)

We used quantitative RT-PCR (qPCR) to examine the expression of marker genes in total RNA isolated from flushed cortical bone from 3 and 12-week-old *Fgfr1^f/f^; Fgfr2^f/f^* and OC-Cre*; Fgfr1^f/f^; Fgfr2^f/f^* mice. Mid-diaphyseal cortical bone was dissected, the marrow was flushed immediately, and the bone tissues were individually frozen in liquid nitrogen and stored at −80°C until analysis. Frozen tissues were pulverized in a dry ice-cooled stainless-steel flask with a ball bearing in a Micro Dismembrator (Sartorius) at 2000rpm for 20 s. RNA was stabilized with Trizol Reagent (#10296**–**010, Life Technologies Corporation, USA) and total RNA isolation was prepared according to the manufacturer’s instructions. cDNA was synthesized using the iScript Select cDNA Synthesis Kit (#1708841, Bio-Rad). mRNA expression was measured using TaqMan Fast Advanced Master Mix (#4444557, Life Technologies) and TaqMan assay probes for *Alp, Axin2, Col1a1, Dkk1, Dmp1, Fgfr1, Fgfr2, Fgfr3, Fgf23, Lef1, Lrp5,* and *Sost. Hprt* was used as a normalization control.

### Sample size and statistics

Based on the coefficient of variation from previous microCT data and a goal to detect a difference in cortical area of 30% (CV = 18%), group sample sizes (n) of n = 6 mice were chosen. For quantitative analysis of cell number, cell death, proliferation, or level of gene expression our goal was to detect a 20% difference between control and experimental samples; a sample size of n = 4-6 mice were chosen based on average CV (∼20%) from recent data. Two-factor ANOVA for DXA, microCT, and histological (osteocyte) measurements; time and genotype were independent factors. Mechanical testing, dynamic histomorphometry, and gene expression measures were evaluated with unpaired t-tests between genotypes groups. All statistical analyses were performed using commercial software (GraphPad Prism 7, La Jolla, CA). Data are represented as mean ± standard deviation (SD). Statistical significance was considered at p < 0.05. In all cases, sample size (n) represents the number of animals per group, which includes approximately equal numbers of males and females.

## Results

### Increased bone formation in mice lacking Fgfr1 and Fgfr2 in mature osteoblasts

To investigate the role of FGFR1 and FGFR2 signaling in mature osteoblasts and osteocytes, *Fgfr1^f/f^; Fgfr2^f/f^* (Fgfr1/2 DFF) mice were crossed with mice carrying the Osteocalcin-Cre (OC-Cre) transgenic allele to generate *Fgfr1*; *Fgfr2* double conditional knockouts (OC-Fgfr1/2 DCKO).^(42–44)^ The OC-Cre transgene becomes active beginning at ∼E17,^(44)^ and targets the mature osteoblast lineage (endosteum, periosteum, and osteocytes). Activity is not detected in non-skeletal tissues.^(44)^ The activity of OC-Cre was confirmed by generating OC-Cre; *ROSA^Tomato^*(Ai9 allele) and OC-Cre; *ROSA^mTmG^*mice. Cre-dependent reporter expression was only observed in the endosteum, periosteum, and osteocytes (Supplemental Figure 1A). Quantitative RT-PCR analysis of mRNA made from flushed cortical bone showed significantly decreased expression of *Fgfr1* and *Fgfr2* in OC-Fgfr1/2 DCKO mice compared to Fgfr1/2 DFF controls, confirming efficient deletion (Supplemental Figure 1B).

Fgfr1/2 DFF control mice and OC-Fgfr1/2 DCKO mice are healthy and fertile and appeared phenotypically normal. Body weight of all mice in the study increased over time, with no differences between OC-Fgfr1/2 DCKO and Fgfr1/2 DFF mice at any timepoint (Figure 1A). *In vivo* DXA imaging showed that the percent body fat remained consistent from 6 to 24 weeks and was not affected by genotype (Figure 1B). However, DXA imaging revealed that whole body bone mineral content (BMC) and areal bone mineral density (aBMD) were both greater in OC-Fgfr1/2 DCKO mice compared to controls, becoming statistically significant at 18 weeks of age (Figure 1C-D). By 24 weeks of age, BMC was 80% higher and aBMD was 30% higher in OC-Fgfr1/2 DCKO mice, with no apparent difference between male and female mice.

**Figure 1.**
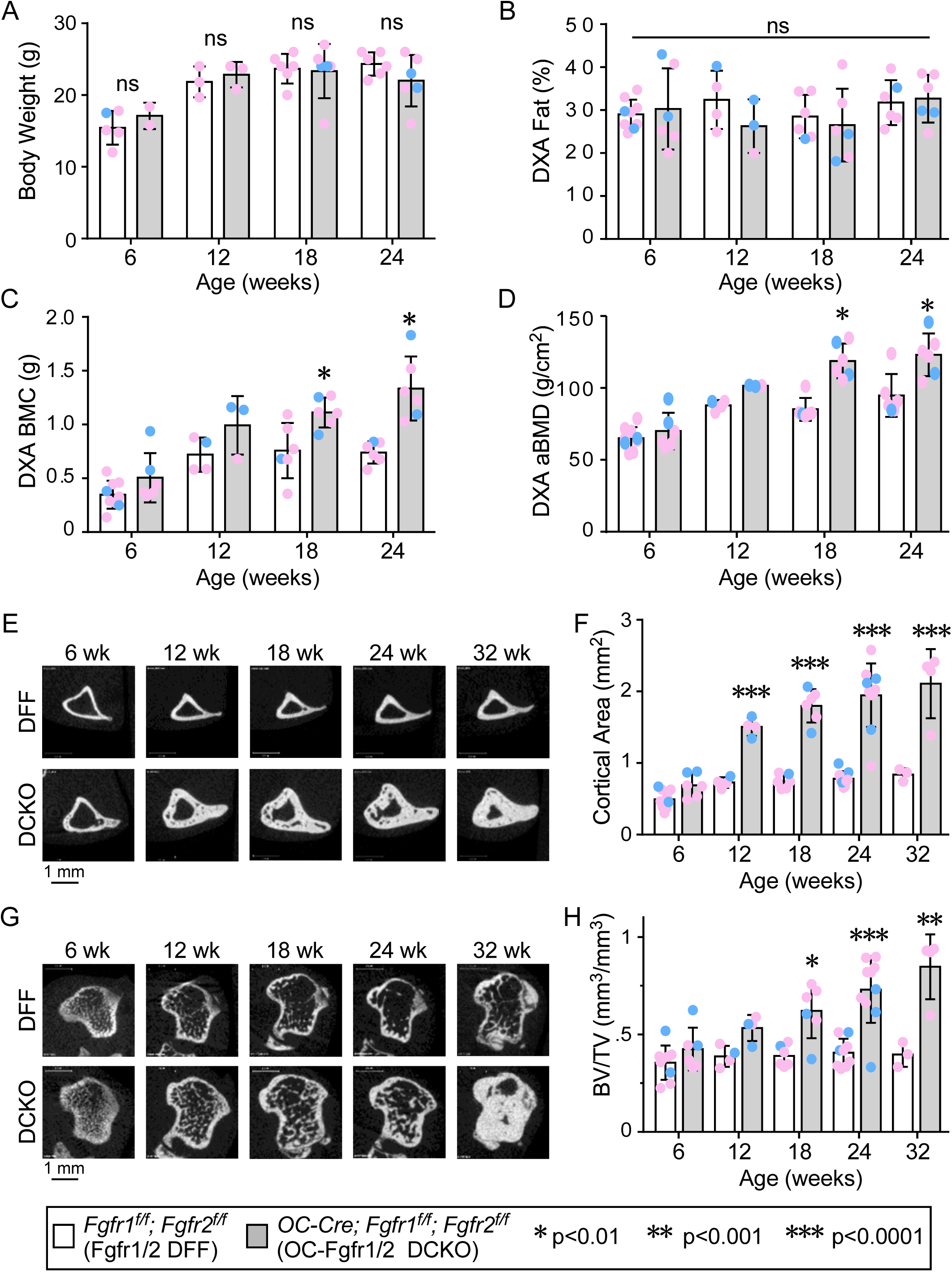
Increased bone mass in mice lacking FGFR1/2 in mature osteoblasts and osteocytes. (A) Body weight measurements showing no effect of genotype between 6 and 24 weeks of age. (B-D) Analysis of whole body (minus head) bone density and body fat content of OC-Fgfr1/2 DCKO mice compared to Fgfr1/2 DFF controls (age 6-24 weeks, n = 6) using dual-energy X-ray absorptiometry (DXA). (B) Measurement of body fat content showing no effect of genotype between 6 and 24 weeks of age. (C, D) Measurement of bone mineral content (BMC) and areal bone mineral density (aBMD) showing a significant increase in OC-Fgfr1/2 DCKO mice by 18 weeks of age. (E-H) Analysis of diaphyseal (cortical) and metaphyseal (cortical + cancellous) bone by in vivo microCT. MicroCT images of mid tibial (E) and proximal tibial (G) sections of Fgfr1/2 DFF mice and OC-Fgfr1/2 DCKO mice. (F) Quantitation of cortical area from mid tibial sections. (H) Quantitation of metaphyseal bone volume/total volume ratio (BV/TV) from proximal tibial sections. Statistics, 2-way ANOVA with Bonferroni’s multiple comparison test. ns, not significant; Pink data points, female mice; Blue data points, male mice.

To quantitatively assess changes in bone structure over time, control and OC-Fgfr1/2 DCKO mice were monitored by *in vivo* microCT between 6 and 32 weeks of age. Compared to controls, OC-Fgfr1/2 DCKO mice revealed a progressive accrual of cortical and cancellous bone occurring after 6 weeks of age in both male and female mice (Figure 1E-H). In the diaphysis of the bone there were no significant differences in cortical area at 6 weeks of age, but by 12 weeks OC-Fgfr1/2 DCKO mice had 107% greater cortical area (Fig 1E-F). Furthermore, total area and pMOI were increased compared to age matched controls in mice 12 weeks and older (Supplemental Table S1). From 18-32 weeks of age there were dramatic increases in total area, cortical area, pMOI, and cortical thickness. Lower cortical TMD (∼8% reduction) was observed in 12-week-old OC-Fgfr1/2 DCKO mice compared to Fgfr1/2 DFF controls. Similar changes were observed in the femur (not shown). Overall, the shape of the long bones appeared normal over this time course. Consistent with changes in diaphyseal cortical bone, the tibia metaphyseal compartment also showed greater TV, BV, BV/TV and vBMD in OC-Fgfr1/2 DCKO mice vs. Fgfr1/2 DFF controls; these differences became significant starting at 18 weeks of age and remained elevated through 32 weeks of age (Figure 1G-H, Supplemental Table S1). By 32 weeks of age, BV was 180% greater in OC-Fgfr1/2 DCKO mice. The results were similar for the metaphyseal region in the distal femur (Supplemental Table S1); however, the femur was more severely affected than the tibia at the 6- and 12-week time points, possibly due to a greater contribution of cortical bone to the metaphyseal measurements in the femur compared to the tibia.

### Increased whole-bone strength but reduced material properties in *OC*-Fgfr1/2 *DCKO* bone

The increased bone mass in OC-Fgfr1/2 DCKO mice suggested that their bones would have increased strength at the whole bone (structural) level. The whole-bone mechanical properties and material properties of femurs from 24-week-old mice were assessed by three-point bending and reference point indentation (RPI). Three-point bending analysis revealed several significant differences in structural mechanical properties of OC-Fgfr1/2 DCKO mice. Compared to controls, OC-Fgfr1/2 DCKO mice showed an 88% increase in yield load, 82% increase in maximum load, and 46% increase in stiffness, but a 71% decrease in post-yield displacement (Figure 2A-D, Supplemental Table S2). Work-to-fracture was unaffected by genotype, due the offsetting effects of increased maximum load and decreased post-yield displacement. MicroCT imaging of the femur midshaft showed increased total area and bone area, and an apparent increased porosity in OC-Fgfr1/2 DCKO mice compared to Fgfr1/2 DFF control mice (Figure 2E). These features resulted in a 5-fold increase in polar moment of inertia (pMOI) (Figure 2H). Estimated material properties, including Young’s modulus, yield stress, and maximum stress, were significantly less by 70%, 44%, and 45%, respectively, in DCKO versus controls (Figure 2F, G, Supplemental Table S2). Reference Point Indentation (RPI) testing demonstrated no significant differences in ID 1^st^, TID, IDI, US 1^st^ or CID parameters between OC-Fgfr1/2 DCKO and Fgfr1/2 DFF mice (Supplemental Table S2).

**Figure 2.**
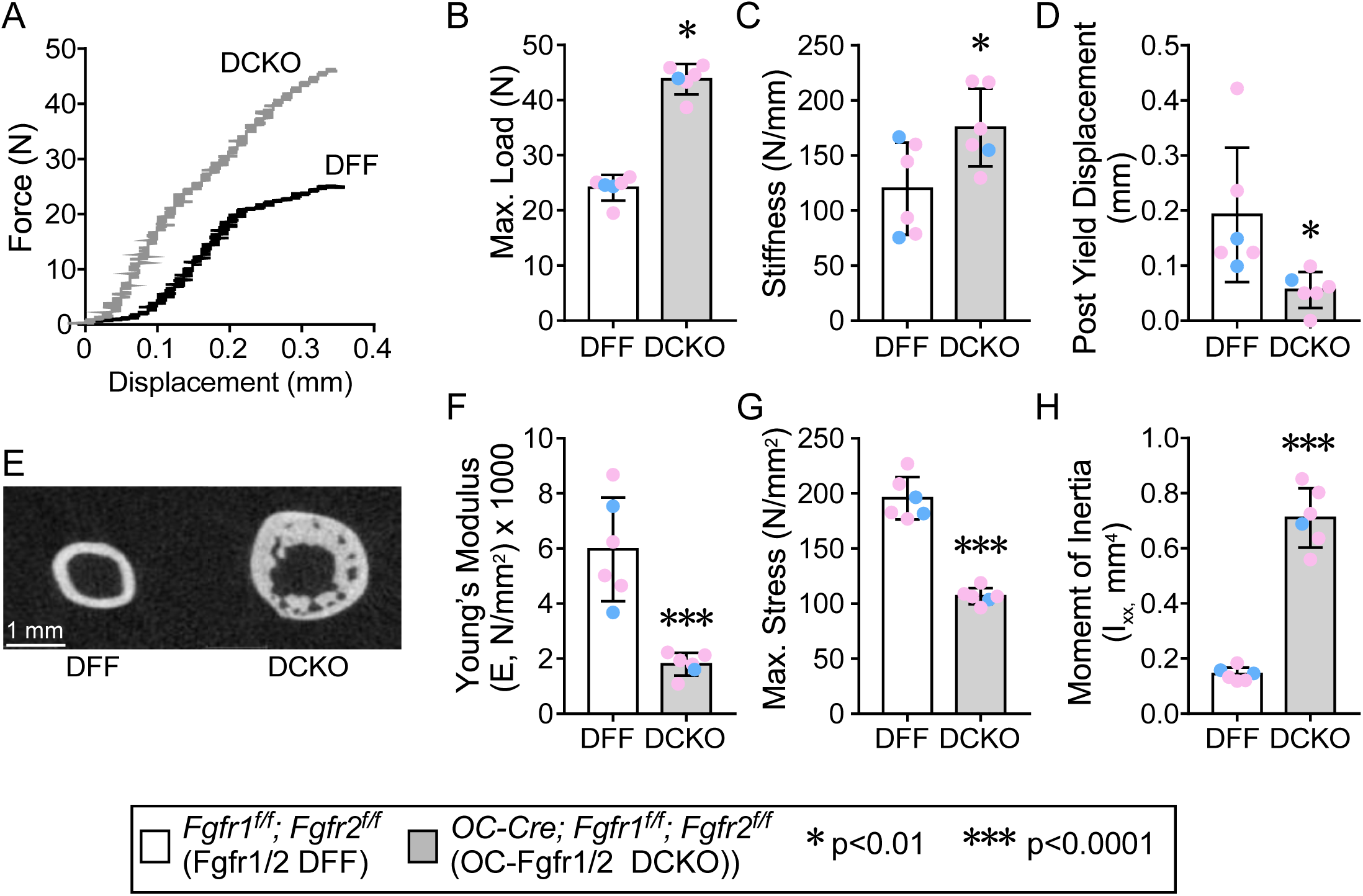
Biomechanical properties of the femurs of OC-Fgfr1/2 DCKO and Fgfr1/2 DFF mice. (A) Femurs from 24-week-old mice were subjected to three-point bending to obtain a force-displacement curve. (B-D) Analysis of these data showing significant differences in maximum load (B), stiffness (C) and post yield displacement (D). (E) MicroCT imaging of the mid-femur showing increased cortical area and porosity in OC-Fgfr1/2 DCKO compared to Fgfr1/2 DFF mice. Analysis of material properties showing a significant decrease in Young’s modulus (F), and maximum stress (G) in OC-Fgfr1/2 DCKO compared to Fgfr1/2 DFF mice. (H) Calculation of the polar moment of inertia showing a significant increase in OC-Fgfr1/2 DCKO compared to Fgfr1/2 DFF mice. Statistics, unpaired Student’s t-test. Pink data points, female mice; Blue data points, male mice.

### Increased mineralized bone and increased osteoblast and osteoclast activity in OC-FGFR1/2 *DCKO* mice

Histological analysis of 12-week-old mice is consistent with the microCT data, showing increased cancellous and cortical bone in OC-Fgfr1/2 DCKO mice compared to Fgfr1/2 DFF control mice (Figure 3A). In contrast, the growth plate showed histologically normal proliferating and hypertrophic chondrocytes (Figure 3B). Cortical bone histology of OC-*Fgfr1/2* DCKO mice shows relatively normal appearing lamellar bone on the periosteal side but an increase in woven bone on the endocortical side (Figure 3A). Mineralization, assessed by Von Kossa staining, showed complete mineralization of all cortical and cancellous regions (Figure 3C). To assess bone structure, histological sections were viewed using polarized light microscopy. Cortical bone from Fgfr1/2 DFF mice showed normal lamellar structure with ordered collagen fibers (Figure 3D), whereas OC-*Fgfr1/2* DCKO mice showed disordered collagen fibers in both lamellar and woven bone regions (Figure 3E).

**Figure 3.**
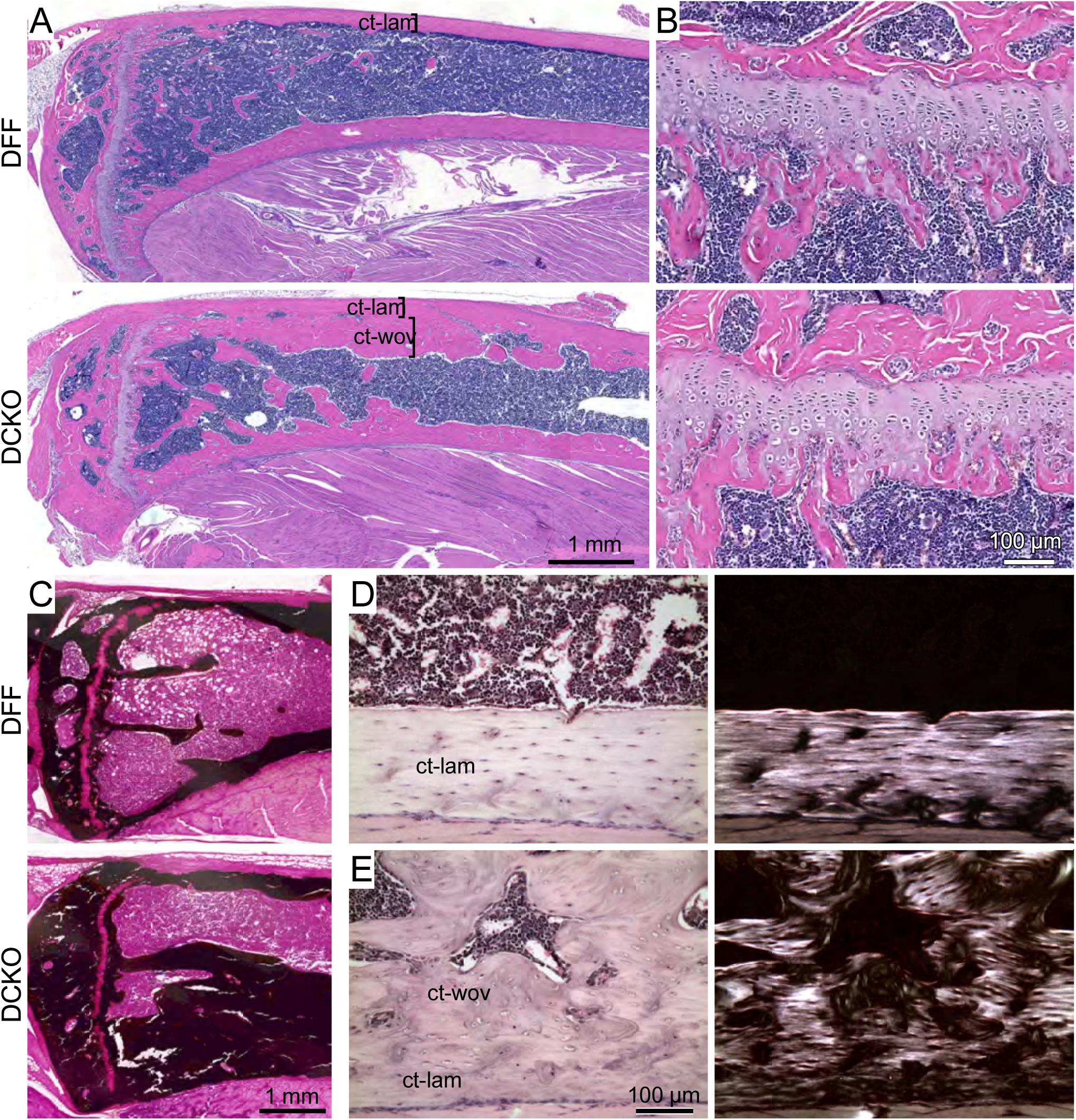
Histological analysis of 12-week-old OC-Fgfr1/2 DCKO and Fgfr1/2 DFF mice. (A) H&E stained sections of the proximal tibia showing increased cortical bone in OC-Fgfr1/2 DCKO compared to Fgfr1/2 DFF mice. (B) Higher magnification view of the growth plate showing normal cytoarchitecture in OC-Fgfr1/2 DCKO and Fgfr1/2 DFF mice. (C) Von Kossa stained sections of the proximal tibia showing increased mineralized bone area in OC-Fgfr1/2 DCKO compared to Fgfr1/2 DFF mice. (D, E) Polarized light microscopy. Bright field compensated images (left) and corresponding polarized light images (right) showing aligned collagen fibers in Fgfr1/2 DFF control cortical bone (D) and disorganized collagen fibers in OC-Fgfr1/2 DCKO lamellar and woven bone (E). ct-lam, cortical lamellar; ct-wov, cortical woven

Dynamic histomorphometry of 12-week-old mice indicated increased anabolic activity in OC-Fgfr1/2 DCKO mice compared to Fgfr1/2 DFF control mice (Figure 4A-J). OC-Fgfr1/2 DCKO mice had a significant increase in endocortical mineral apposition rate (MAR) (61%) and bone formation rate (BFR/BS) (55%), with no changes in mineralizing surface (MS/BS). On the periosteal surface, MAR, MS/BS and BFR/BS were all significantly increased by 35-164%. In addition, abundant double label is visible within the cortical bone of the OC-Fgfr1/2 DCKO, which is not present in the Fgfr1/2 DFF mouse.

**Figure 4.**
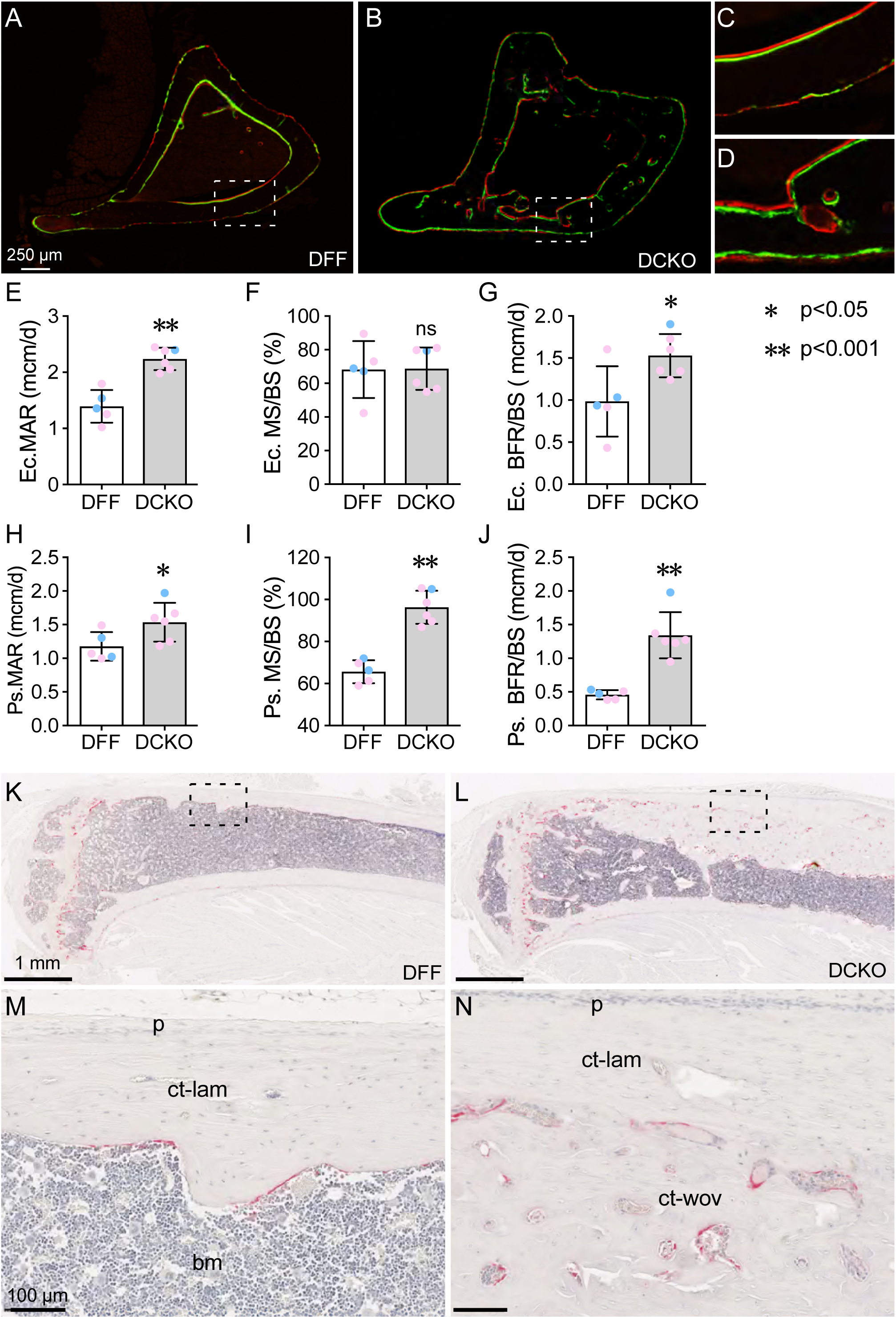
Increased bone growth and remodeling in mice lacking FGFR1/2 in mature osteoblasts and osteocytes. (A-J) Calcein and alizarin labeled transverse sections from the tibial diaphysis of Fgfr1/2 DFF (A), OC-Fgfr1/2 DCKO (B) mice at 12 weeks of age. (C, D) shows magnified views for the boxed regions in A and B, respectively. Image analysis of endocortical bone shows an increased mineral apposition rate (E) and bone formation rate/bone surface ratio (G), but no difference in the mineralizing surface/bone surface ratio (F) in OC-Fgfr1/2 DCKO compared to Fgfr1/2 DFF mice. Image analysis of periosteal bone shows an increased mineral apposition rate (H), mineralizing surface/bone surface ratio (I), and bone formation rate/bone surface ratio (J) in OC-Fgfr1/2 DCKO compared to Fgfr1/2 DFF mice. (K-N) TRAP staining at 12 weeks of age shows increased numbers of osteoclasts in cortical bone in OC-Fgfr1/2 DCKO (L, N) compared to Fgfr1/2 DFF (K, M) mice. ct-lam, cortical lamellar; ct-wov, cortical woven; bm, bone marrow; p, periosteum; ns, not significant. Statistics, unpaired Student’s t-test. Pink data points, female mice; Blue data points, male mice.

Both the histology and dynamic histomorphometry indicate an increase in the porosity of the cortical bone in OC-Fgfr1/2 DCKO mice, primarily in the endocortical woven bone region (Figure 4B, D, L, N). TRAP staining, used to identify osteoclasts, shows that cortical porosities in OC-Fgfr1/2 DCKO mice are lined with osteoclasts, indicating active cortical remodeling (Figure 4K-N).

### Osteocyte loss precedes increased bone accrual

A striking histological feature of OC-Fgfr1/2 DCKO bone histology is the absence of nuclei within osteocyte lacunae, observed in cortical bone (Figure 5), trabecular bone, and vertebral bodies (not shown). Osteocyte lacunae were histologically scored as alive, dying, or dead, based on lacunar morphology (Figure 5J). At 3 weeks of age, OC-Fgfr1/2 DCKO cortical bone showed normal lamellar histology (Figure 5A, B). However, quantitation of nucleated lacunae (classified as alive cells) showed a 36% decrease in OC-Fgfr1/2 DCKO tibia compared to Fgfr1/2 DFF control tibia (p < 0.01) and a corresponding increase in dying/dead osteocytes (Figure 5A-C). TUNEL staining at 3 weeks of age showed a similar proportion of labeled apoptotic cells (Figure 5M, N). Consistent with normal appearing cortical bone, TRAP staining of 3-week-old OC-Fgfr1/2 DCKO mice did not reveal qualitative differences in the abundance or appearance of osteoclasts (Supplemental Figure 2). At 6 weeks of age cortical histology also showed normal appearing lamellar bone; however, there was a 64% decrease in alive osteocytes (p < 0.001) and a corresponding increase in dying/dead osteocytes (Figure 5D-F).

**Figure 5.**
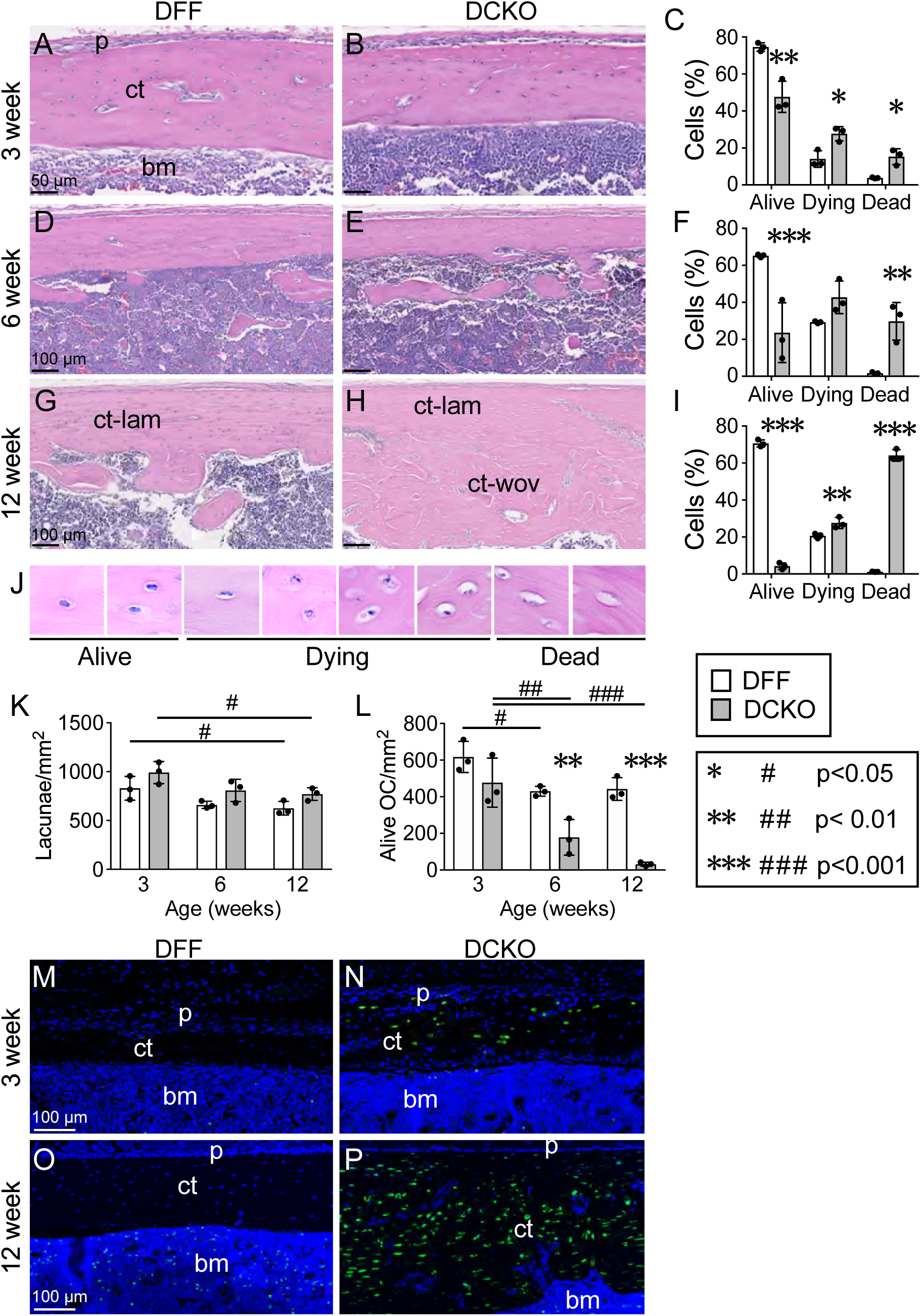
Osteocyte loss in mice lacking FGFR1/2 in mature osteoblasts and osteocytes. (A-L) Analysis of osteocyte lacunae morphology at 3, 6, and 12 weeks of age. Osteocytes were categorized according to the morphologies shown in panel J. Three-week-old mice (A-C) showing decreased numbers of “alive” osteocytes and increased numbers of “dead” and “dying” osteocytes. These differences were progressively increased at 6 weeks of age (D-F) and 12 weeks of age (G-I). (K) Analysis of lacunar density showing no effect of genotype at any age. (L) Analysis of “alive” osteocyte density showing no difference at 3 weeks of age, but reduced density at 6 and 12 weeks of age. (M-P) TUNEL staining showing increased osteocyte apoptosis in cortical bone from 3-week-old (M, N) and 12-week-old (O, P) OC-Fgfr1/2 DCKO compared to Fgfr1/2 DFF mice. ct-lam, cortical lamellar; ct-wov, cortical woven; bm, bone marrow; p, periosteum. Statistics: Alive, dying, dead, cells, OC density analyzed by 2-way ANOVA with Sidak’s multiple comparison test; Lacunae density analyzed by 2-way ANOVA with Tukey’s multiple comparison test.

Twelve-week-old mice showed a layer of normal-appearing lamellar bone on the periosteal side, and a large increase in endocortical woven bone. In both histological domains there was a dramatic absence of alive osteocytes (94% reduction, p < 0.001) in OC-Fgfr1/2 DCKO mice compared to Fgfr1/2 DFF controls and a large (54 fold) increase in the number of dead osteocyte lacunae (Figure 5G-I). The fragmented appearing osteocyte nuclei suggested apoptosis, which was confirmed by TUNEL assay, which showed abundant staining in OC-Fgfr1/2 DCKO cortical bone, but not in Fgfr1/2 DFF control bone (Figure 5O, P). At all three ages examined, the overall density of osteocyte lacunae remained similar in Fgfr1/2 DFF and OC-Fgfr1/2 DCKO mice (Figure 5K). Both genotypes showed a moderate (22-24%, p < 0.05) decrease in lacunar density between 3 and 12 weeks of age. Interestingly, at 3 weeks of age, the average density of alive osteocytes in OC-Fgfr1/2 DCKO mice was similar to that of controls at all time points (Figure 5L). This suggests that the density of living osteocytes may be the limiting factor for normal cortical bone growth.

### Inactivation of FGFR1 is responsible for osteocyte loss and high bone mass

To assess the relative contributions of *Fgfr1* and *Fgfr2* to the regulation of bone growth, we generated OC-Cre; *Fgfr1^f/f^; Fgfr2^f/+^* and OC-Cre; *Fgfr1^f/+^; Fgfr2^f/f^* single conditional knockout (CKO) mice that are heterozygous for a floxed allele of the other *Fgfr*. Analysis of these mice at 6, 12 and 18 weeks of age by *in vivo* microCT showed that the increased bone mass phenotype in OC-Fgfr1/2 DCKO mice can be accounted for by inactivation of *Fgfr1* alone (Figure 6, Supplemental Table S3). Furthermore, OC-Cre; *Fgfr1^f/+^; Fgfr2^f/f^* mice also serve as controls for OC-Cre, showing that the OC-Cre allele does not contribute to any of the noted bone phenotypes and that one wild type allele of *Fgfr1* is sufficient for normal bone growth. Histological analysis of tibia from 12-week-old mice shows normal cortical bone and osteocytes in OC-Cre; *Fgfr1^f/+^; Fgfr2^f/f^* mice and increased cortical thickness and osteocyte loss in OC-Cre; *Fgfr1^f/f^; Fgfr2^f/+^* mice (Figure 6C, D).

**Figure 6.**
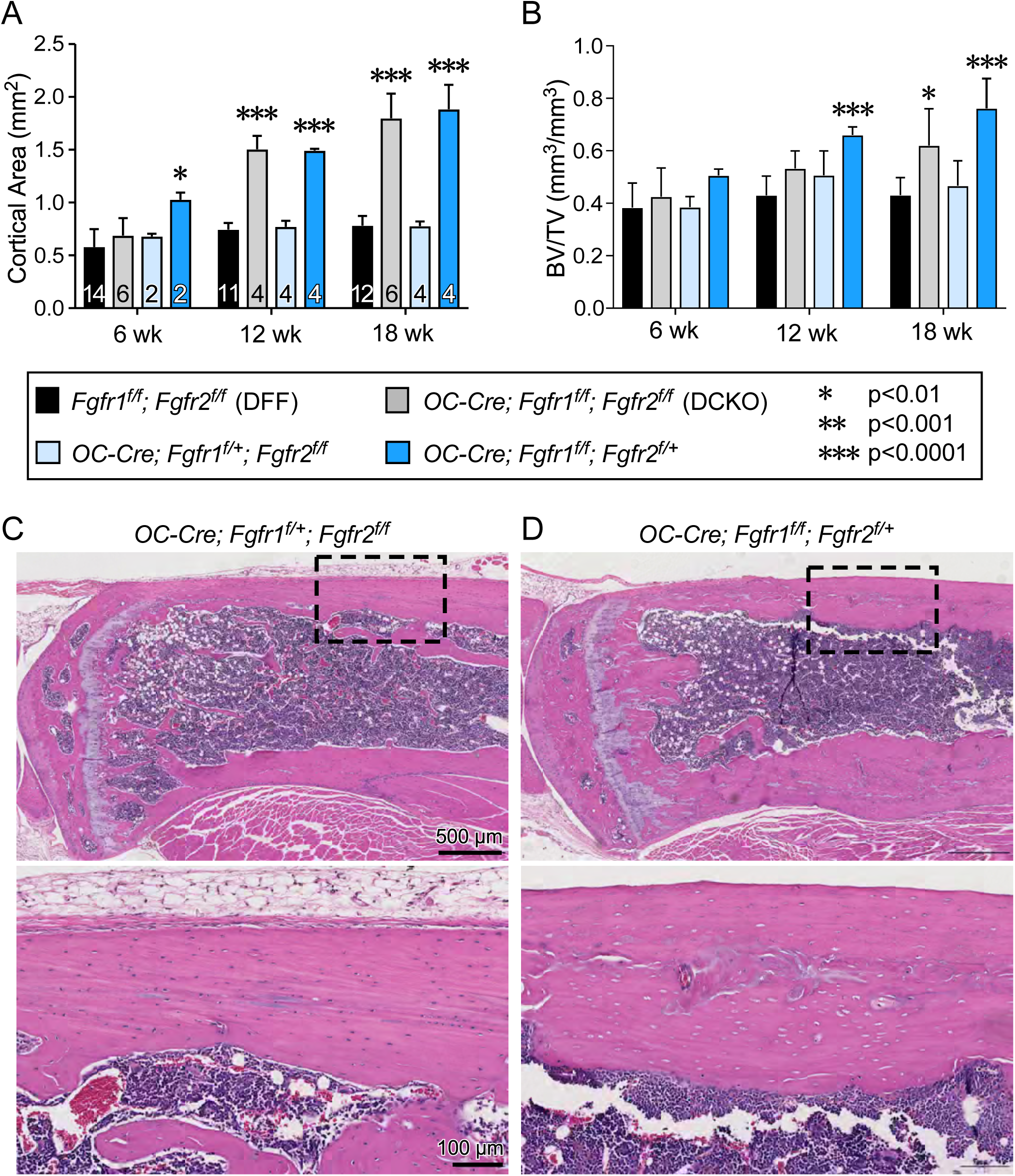
Analysis of cortical bone in different allelic combinations of *Fgfr1* and *Fgfr2* targeted with OC-Cre demonstrating that increased bone mass only occurs when both copies of *Fgfr1* are deleted. (A) Quantitation of cortical area from mid tibial sections. (B) Quantitation of bone volume/total volume ratio (BV/TV) from proximal tibial sections. Statistics, 2-way ANOVA; group sample sized indicated by number within each bar. (C) Representative histological sections from mice lacking *Fgfr2* and heterozygous for *Fgfr1*. (D) Representative histological sections from mice lacking *Fgfr1* and heterozygous for *Fgfr2*. Lower panels show higher magnification of the boxed regions above.

### Increased bone formation correlates with activation of Wnt/βCatenin signaling

Gene expression was analyzed in cortical bone in 3- and 12-week-old mice by quantitative RT-PCR (Figure 7). At 3 weeks of age, markers of bone formation and osteoblast maturation, *Col1a1*, *Alp*, and *Fgfr3*, were decreased in OC-Fgfr1/2 DCKO mice compared to Fgfr1/2 DFF control mice, suggesting that an initial consequence of loss of FGFR1 and FGFR2 was decreased bone formation. This is consistent with reduced Wnt/βCatenin signaling, shown by lower levels of *Axin2*, *Lrp5,* and *Lef1*. Consistent with the initiation of osteocyte loss, levels of *Dmp1*, *Sost,* and *Dkk1,* were decreased, but paradoxically, levels of *Fgf23* were increased. At this stage, clearly the decreased level of *Sost* and *Dkk1* does not correlate with reduced Wnt/βCatenin signaling and increased level of *Fgf23* does not correlate with osteocyte loss. At 12 weeks of age, when robust woven bone formation was observed in OC-Fgfr1/2 DCKO mice, markers of bone formation, *Col1a1*, *Alp*, and *Fgfr3*, were increased. Consistent with loss of most osteocytes and concurrent increased bone formation, *Sost* expression was decreased and markers of Wnt/βCatenin signaling, *Lrp5* and *Lef1,* were increased in OC-*Fgfr1/2* DCKO mice compared to *Fgfr1/2* DFF control mice. Notably, in 12-week-old mice, levels of *Dkk1* were increased, possibly as a mechanism to compensate for loss of Sost.^(13)^ Similar to 3-week-old OC-Fgfr1/2 DCKO mice, levels of *Fgf23* remained high in 12-week-old mice, despite the decreased number of live osteocytes. Finally, immunostaining also showed fewer Sclerostin and DMP1 positive cells within cortical bone of OC-Fgfr1/2 DCKO mice (Supplemental Figure 3).

**Figure 7.**
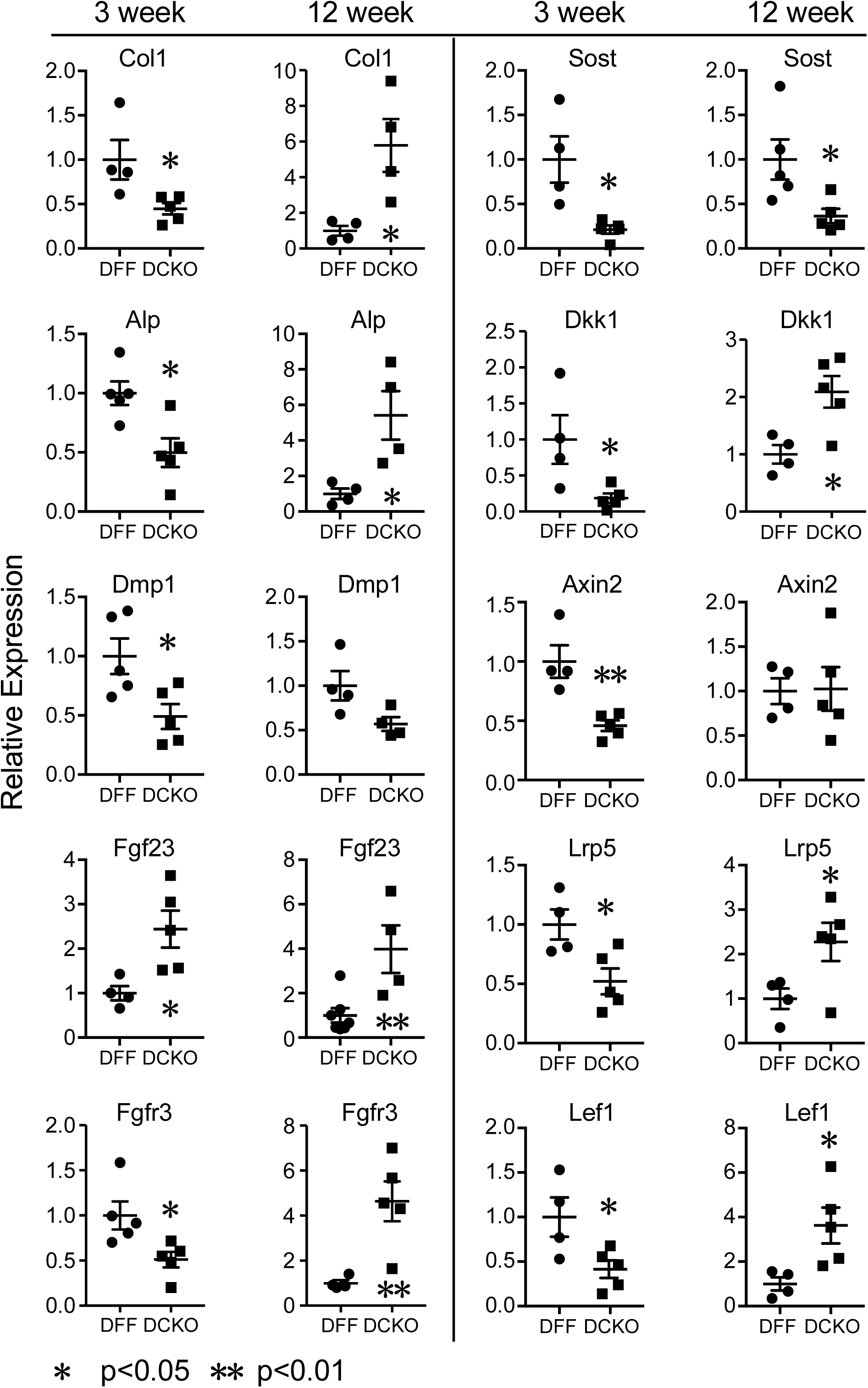
Gene expression in cortical bone from OC-Fgfr1/2 DCKO and Fgfr1/2 DFF mice. Cortical bone (femur and tibia) from 3- and 12-week-old mice were flushed to remove bone marrow and used to prepare total RNA. Quantitative RT-PCR was used to examine changes in gene expression of osteoblast anabolic bone genes (*Col1a1, ALP, Fgfr3*), genes preferentially expressed in osteocytes (*Dmp1, Fgf23, Sost, Dkk1*), and indicators of Wnt/β-catenin signaling activity (*Axin2, Lrp5, Lef1*). Statistics, unpaired Student’s t-test.

### FGFR requirements for osteocyte survival in mature mice

The phenotype caused by inactivation of *Fgfr1* and *Fgfr2* with OC-Cre may be due to altered development in the late embryonic or neonatal period, and/or to altered FGFR1/2 signaling in later postnatal osteoblasts and osteocytes. To distinguish between these possibilities, we used the Dmp1-CreER allele to initiate the inactivation of *Fgfr1* and *Fgfr2* beginning in 3-week-old mice. To compare cell populations targeted by Dmp1-CreER and OC-Cre, lineage reporters were used. Lineage analysis of Dmp1-CreER; *ROSA26^Tomato^*(Ai9 allele) mice, induced with tamoxifen beginning at 3 weeks of age for 2 weeks, is very similar to that of adult OC-Cre, *ROSA26^Tomato^* mice, with expression in both osteoblasts and osteocytes (compare Figure 8A, B and Supplemental Figure 1).

**Figure 8.**
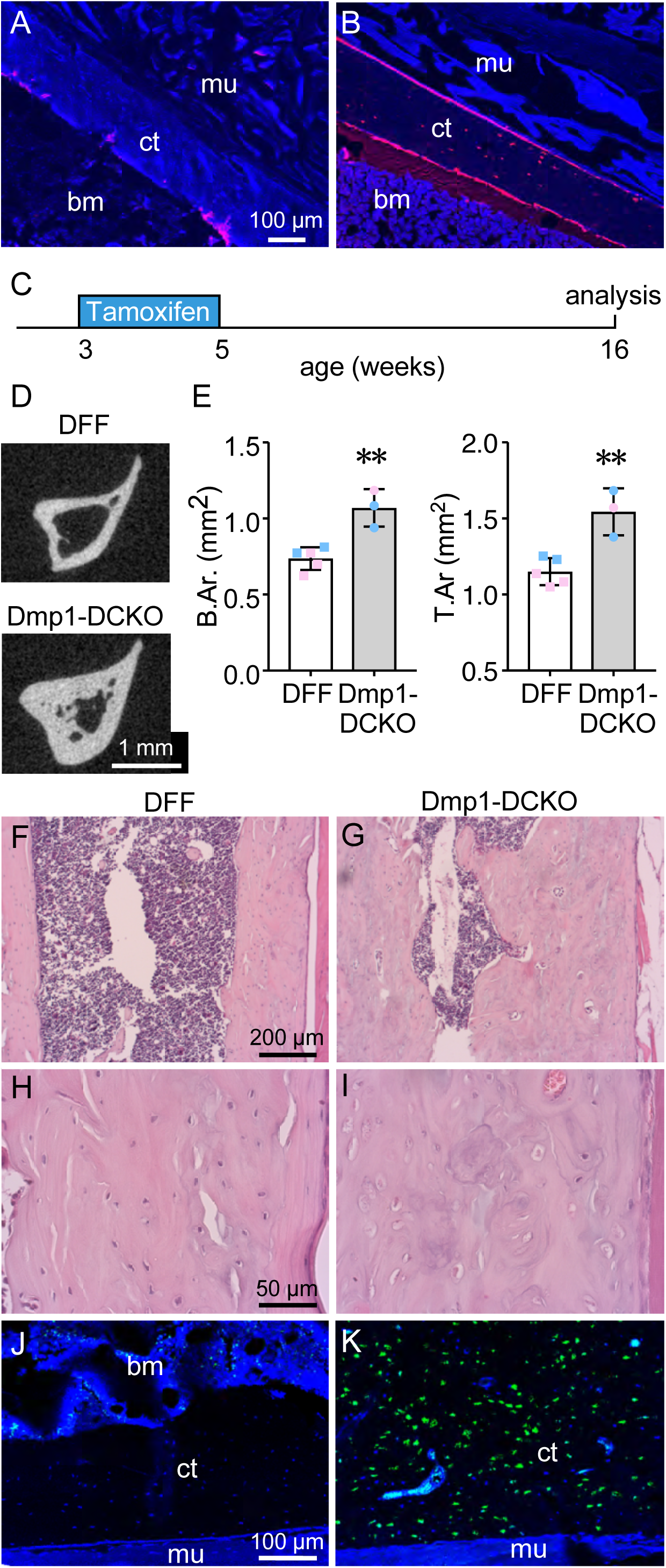
Localization of the Dmp1-CreER lineage and inactivation of *Fgfr1* and *Fgfr2* in mature osteoblasts and osteocytes in postnatal mice. (A, B) Fluorescence imaging for Tomato (red) in cortical bone from *ROSA26^Tomato^*mice (A) and Dmp1-CreER*; ROSA26^Tomato^* mice (B) that were induced with tamoxifen chow from 3-5 weeks of age and analyzed at 6 weeks of age. (C) Experimental timeline for conditional knockout of *Fgfr1* and *Fgfr2* in postnatal mature osteoblasts and osteocytes. (D) MicroCT imaging of the mid-tibia in 16-week-old mice showing increased cortical area and porosity in tamoxifen induced Dmp1-Fgfr1/2 DCKO compared to Fgfr1/2 DFF control mice. (E) Quantitation of cortical bone area and total area. (F-I) Histology (H&E stain) showing mid-tibia osteocyte lacunae morphology in tamoxifen-induced Dmp1-Fgfr1/2 DCKO and Fgfr1/2 DFF control mice at 16 weeks of age. Low magnification images (F, G) and high magnification images (H, I) are shown. (J, K) TUNEL staining showing increased osteocyte apoptosis in cortical bone from 16-week-old Dmp1-Fgfr1/2 DCKO compared to Fgfr1/2 DFF control mice that were induced with tamoxifen from 3-5 weeks of age. bm, bone marrow; ct, cortical bone; mu, muscle. Statistics, unpaired Student’s t-test, ** p < 0.01. Pink data points, female mice; Blue data points, male mice.

Dmp1-CreER; *Fgfr1^f/f^; Fgfr2^f/f^* (DMP-Fgfr1/2 DCKO) mice and Fgfr1/2 DFF control mice were given tamoxifen chow beginning at 3 weeks of age for 2 weeks to inactivate *Fgfr1* and *Fgfr2* in osteoblasts and osteocytes. Mice were then aged to 16 weeks and limbs were examined by microCT followed by histology (Figure 8C). MicroCT analysis of 16-week-old DMP-Fgfr1/2 DCKO mice showed increased bone area and total area in the mid tibia (Figure 8D, E, Supplemental Table S4) and mid femur (not shown). Histological analysis showed increased cortical bone accompanied by osteocyte lacunae that were empty or contained fragmented nuclei (Figure 8F-I). Similar to the OC-Fgfr1/2 DCKO mice, DMP-Fgfr1/2 DCKO mice showed positive TUNEL staining throughout cortical bone (Figure 8J-K).

## Discussion

In this report we identify a novel function of FGFR signaling in mature osteoblasts and osteocytes which is required to maintain osteocyte viability. Inactivation of *Fgfr1* and *Fgfr2* or only *Fgfr1* in late embryonic stages with the OC-Cre allele resulted in osteocyte loss beginning as early as 3 weeks of age. Inactivation of *Fgfr1* and *Fgfr2* beginning at 3 weeks of age using the tamoxifen-inducible Dmp1-CreER allele also led to osteocyte loss, observed at 12 (data not shown) and 16 weeks of age. Because both of these Cre alleles target mature osteoblasts and osteocytes, it is possible that FGFR signaling directly within osteocytes and/or that FGFR signaling in immediate osteocyte precursors or mature osteoblasts is required for osteocyte survival.

Osteocytes are a source of Sclerostin (product of the *Sost* gene) and DKK1, Wnt inhibitory proteins that disrupt interactions of Wnt ligands with the co-receptor LRP5 and LRP6.^(10,51–54)^ Reduced expression of *Sost* and *Dkk1* would be expected to lead to increased Wnt/βCatenin signaling and stimulation of osteoblast anabolic activity.^(10,55)^ However, in 3-week-old OC-Fgfr1/2 DCKO mice, *Sost* and *Dkk1* expression were reduced while Wnt/βCatenin signaling, as assessed by expression of Wnt reporter genes, showed no evidence of activation and in fact were reduced. Thus, at 3 weeks, other factors, such as reduced FGFR1 mediated osteoanabolic activity, may mask effects of reduced expression of Wnt inhibitory proteins. In contrast, by 12 weeks of age, when most osteocytes were lost, *Sost* was also reduced and, as expected, some markers of Wnt/βCatenin signaling (*Lrp5*, *Lef1*) were increased. Increased expression of *Dkk1* at 12 weeks may be secondary to a compensatory effect of reduced *Sost* and increased Wnt signaling, as seen by other investigators.^(56–58)^ We posit that increased Wnt/βCatenin signaling beginning between 6 and 12 weeks of age is one factor that contributes to the observed increase in bone formation observed in 12-week and older mice.

The mechanism(s) by which loss of FGFR signaling results in osteocyte loss is not known and could involve regulation of direct cell survival pathways, regulation of the osteocyte lacunocanalicular network or the surrounding extracellular matrix, or other signaling systems in bone that affect osteocyte survival. Regulation of the lacunocanalicular network is particularly attractive given that through its transduction of mechanical force the lacunocanalicular network is important for osteocyte survival.^(17,59)^ The organization of the lacunocanalicular network correlates with that of the extracellular matrix.^(60)^ Polarizing light microscopy shows that the lamellar bone structure is less ordered in OC-Fgfr1/2 DCKO mice, and this could disrupt the osteocyte lacunocanalicular network leading to osteocyte death. Other factors that promote osteocyte apoptosis include estrogen withdrawal, glucocorticoids, unloading, and microdamage^(17,61,62)^ Although the mechanisms by which these events lead to osteocyte apoptosis are poorly understood, our results raise the possibility that altered FGFR signaling may play a role. Examination of mechanisms by which FGFR signaling promotes osteocyte survival will be the subject of future studies.

The increase in bone mass in OC-Fgfr1/2 DCKO is preceded by the initial loss of osteocytes, suggesting that increased bone accrual is secondary to osteocyte loss. This possibility could be tested by pharmacologically inhibiting osteocyte apoptosis with the pan-caspase inhibitor QVD, which has been shown to block osteocyte apoptosis and prevent pathological bone remodeling.^(17,63–65)^

At both 3 and 12 weeks of age, we observed increased *Fgf23* mRNA expression in cortical bone. Given that the number of viable osteocytes is in decline, this suggests that the increased *Fgf23* expression likely occurs in osteoblasts or pre-osteocytes.^(66)^ Interestingly, it has been shown that viable osteocytes, within 100 to 300 µm from dying osteocytes, have increased expression of VEGF and RANKL.^(62,67)^ We posit that *Fgf23* expression may be similarly increased in nearby viable osteocytes or pre-osteocytes in response to their dying neighbors. The mechanisms leading to increased *Fgf23* in OC-Fgfr1/2 DCKO mice are not known and likely to be indirect since available data suggests that FGFR1 signaling functions to directly promote *Fgf23* expression in osteocytes and bone marrow stromal cells.^(68–70)^ Furthermore, genetic inactivation of *Fgfr1* in osteocytes of Hyp mice, a mouse model of X-linked hypophosphatemia (XLH), reduced the excessive FGF23 levels in osteocytes and partially rescued the bone phenotype in these mice.^(71)^ In contrast, pharmacological activation of FGFR1 in osteoblasts led to increased FGF23 secretion and hypophosphatemia in adult mice.^(72)^ These data support an important role for FGF23/FGFR1 signaling in the control of bone mass and mineralization *in vivo*, but through a mechanism that is distinct from that in OC-Fgfr1/2 DCKO mice which lack FGFR1 in mature osteoblasts and osteocytes.

*Fgfr3* is expressed in mature osteoblasts and osteocytes.^(20,35,36)^ Because most osteocytes are lost in OC-Fgfr1/2 DCKO mice at 12 weeks, it is likely that *Fgfr3* expression is increased in mature osteoblasts or other cell types. Although it is not known if FGFR3 signaling is increased, combination of increased FGF23 and increased FGFR3 provides a potentially active ligand-receptor pair. One function of FGF23-FGFR3 signaling in bone is to suppress tissue non-specific alkaline phosphatase (TNAP) expression and reduce mineralization.^(20,73)^ In the case of OC-Fgfr1/2 DCKO mice, this could be a compensatory response to the massive increase in bone formation at 12 weeks.

The FGF ligands that function in neonatal and juvenile growing bone are not known. FGF2, FGF9, and FGF18 are good candidates since FGF9 and FGF18 have established roles in bone development,^(33,74–81)^ and FGF2 has established roles in growing and mature bone.^(82)^ However, loss of FGF2 leads to osteoporosis, suggesting that it functions at a different stage of the osteoblast lineage, or functions through a different FGFR, such as FGFR3, which is also expressed in mature osteoblasts. Future experiments will be required to determine if there is redundancy between FGF2 and FGF9/18, or unique stage-specific activities of these ligands during osteoblast maturation in growing bone.

In conclusion, these studies establish a role for FGFR signaling in the mature osteoblast lineage in growing bone in neonatal and juvenile mice. Loss of FGFR signaling in mature osteoblasts or osteocytes either directly or indirectly leads to osteocyte death, which in turn leads to increased cortical remodeling and eventual bone mass accrual. Future studies will be required to determine whether FGFR signaling has a homeostatic role in mature bone. Limitations of these studies include efficiency of conditional knockout alleles, possible effects of genetic background, and the range of time points (from 3 to 32 weeks) that were analyzed.

## Acknowledgements

The authors thank D. Leib and C. Idleburg and the Washington University Musculoskeletal Research Center for help with MicroCT and histology, and A. Robling and P. Divieti Pajevic for providing Dmp1-CreER mice.

## Authors’ Roles

Study design: JM, KK, MS and DO. Study conduct: JM, KK and CS. Data collection: JM, KK, CS and JL. Data interpretation: JM, CS, KK, MS and DO. Drafting manuscript: JM and DO. Revising manuscript: JM, CS, KK, MS and DO. Approving final version of manuscript: JM, MS and DO. JM, CS, MS, and DO take responsibility for the integrity of the data analysis.

## Supplemental Data

**Supplemental Figure 1.**
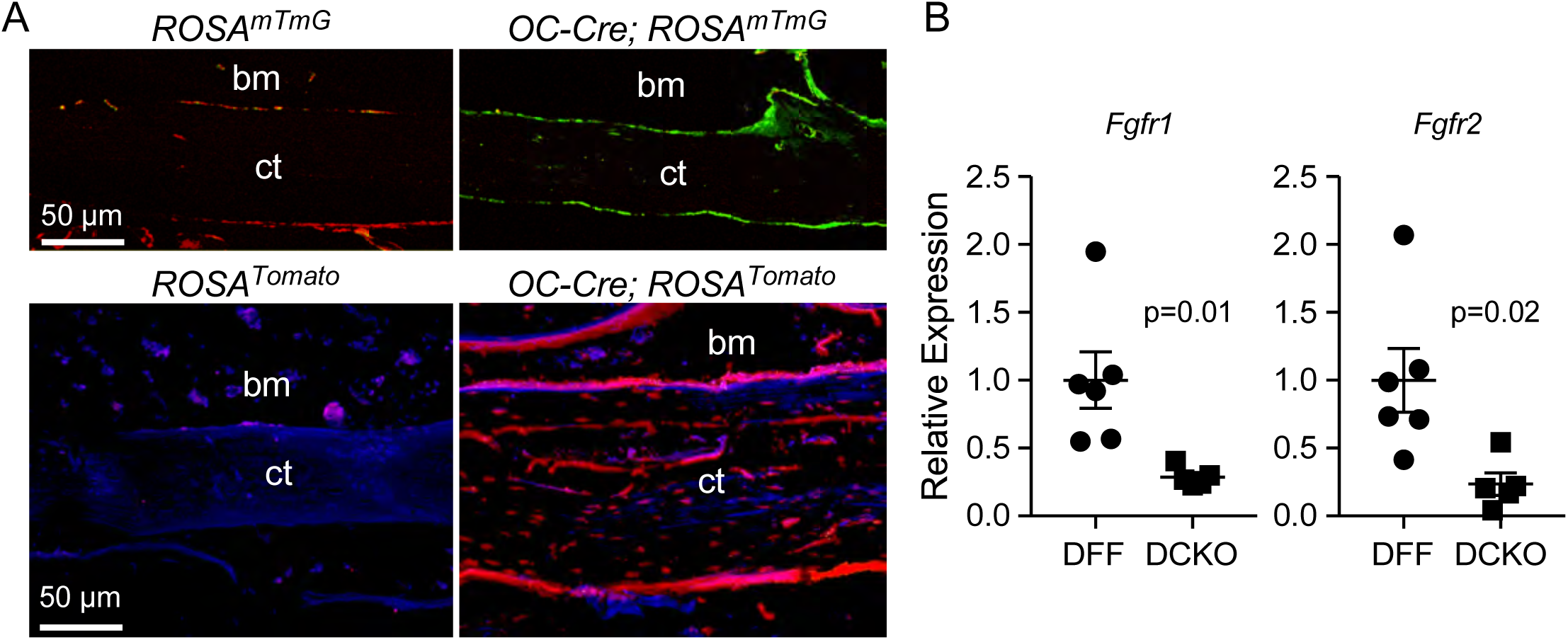
Localization of the OC-Cre lineage and expression of *Fgfr1* and *Fgfr2* following conditional inactivation with OC-Cre. (A) *ROSA^mTmG^* and *ROSA^Tomato^* reporters. Recombination of the *ROSA^mTmG^* allele exposed to OC-Cre activates membrane bound GFP (mG) in osteoblasts and osteocytes. Non-targeted cells express membrane bond Tomato (mT). Recombination of the *ROSA^Tomato^*allele exposed to OC-Cre activates Tomato (red) in osteoblasts and osteocytes. Blue, DAPI; bm, bone marrow; ct, cortical bone. (B) Quantitative RT-PCR at 3 weeks of flushed cortical bone showing reduced expression of *Fgfr1* (3.5 fold) and *Fgfr2* (4.2 fold) in OC-Fgfr1/2 DCKO mice compared to Fgfr1/2 DFF control mice.

**Supplemental Figure 2.**
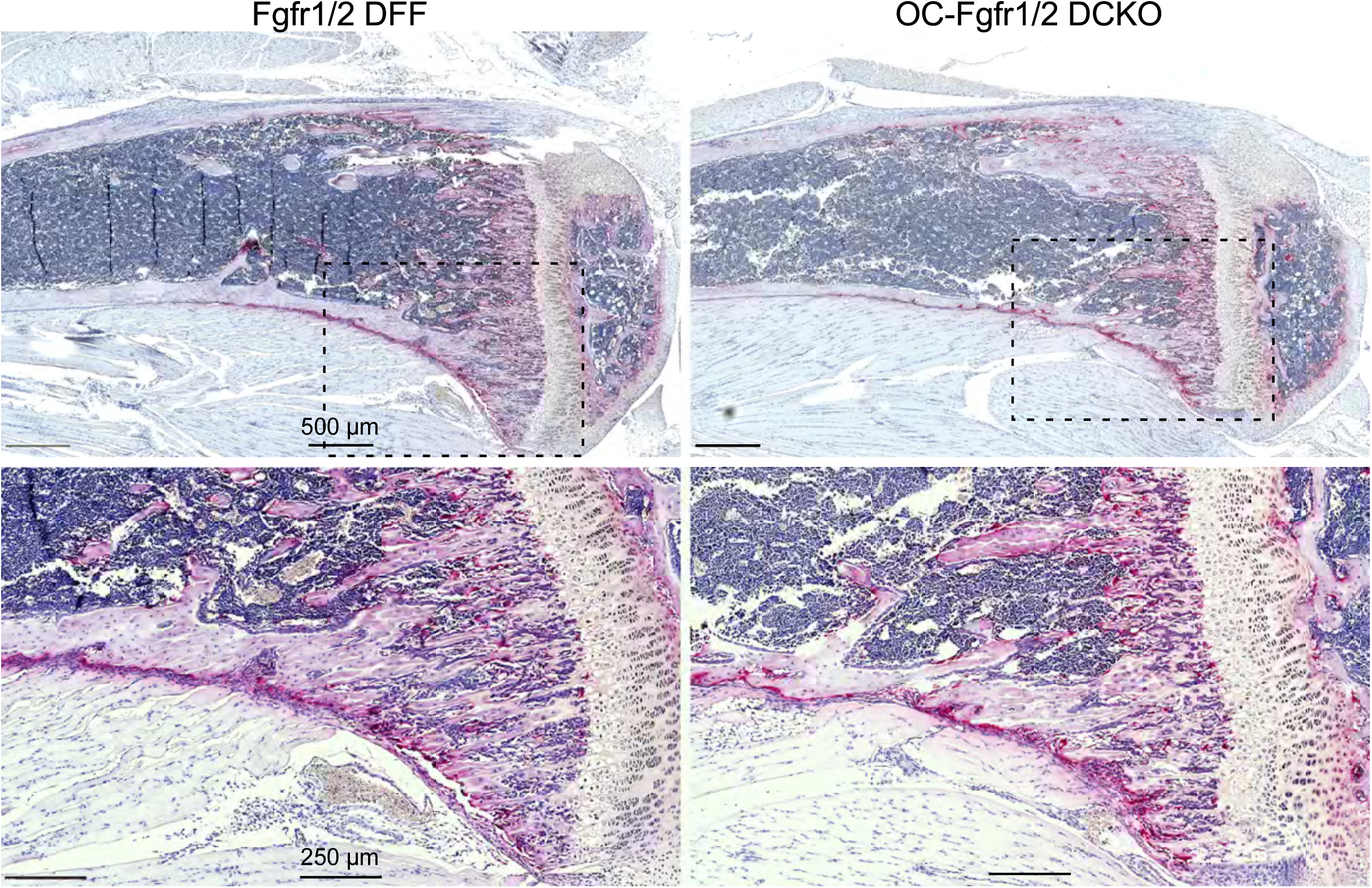
TRAP staining at 3 weeks of age showing similar numbers of osteoclasts in cortical bone and trabecular bone in OC-Fgfr1/2 DCKO compared to Fgfr1/2 DFF mice. Lower panels show higher magnification of the boxed regions above.

**Supplemental Figure 3.**
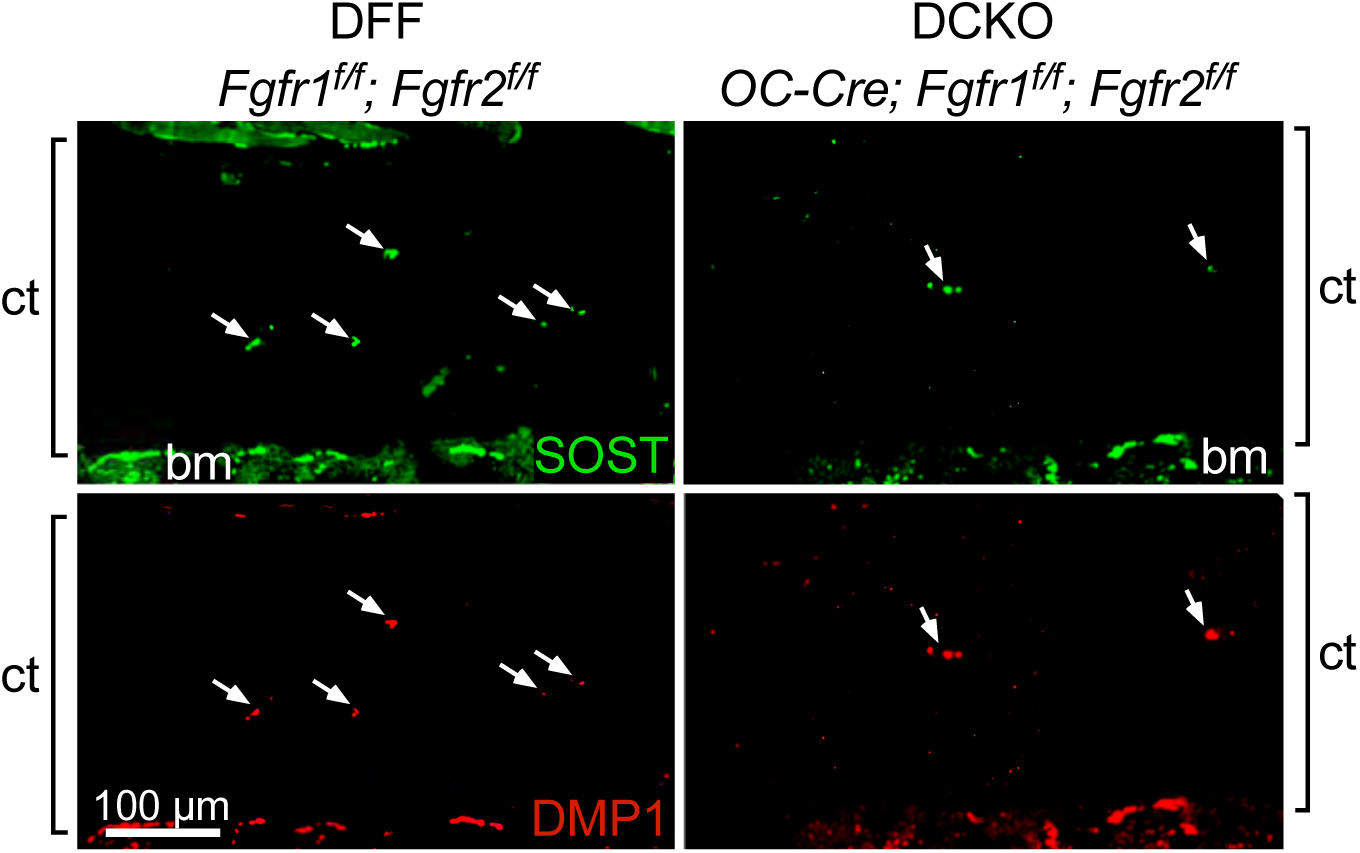
Expression of osteocyte markers, *Sost* and *Dmp1*, in cortical bone from 12-week-old OC-Fgfr1/2 DCKO and Fgfr1/2 DFF mice. bm, bone marrow; ct, cortical bone.

**Supplemental Table S1.** MicroCT analysis of tibia and femur in Fgfr1/2 DFF and OC-Fgfr1/2 DCKO mice

|  | 6 wk |  |  |  | 12 wk |  |  |  |  |  |  |  |
| --- | --- | --- | --- | --- | --- | --- | --- | --- | --- | --- | --- | --- |
|  | #DFF |  | #DCKO |  | DFF |  | DCKO |  |  |  |  |  |
|  | Mean | St.Dev. | Mean | St.Dev. | Mean | St.Dev. | Mean | St.Dev. |  |  |  |  |
| **Tibia Cortical Bone** |  |  |  |  |  |  |  |  |  |  |  |  |
| T.Ar (mm<sup>2</sup>) | 1.45 | 0.20 | 1.73 | 0.07 | 1.22 | 0.17 | 2.15*** | 0.19 |  |  |  |  |
| B.Ar (mm<sup>2</sup>) | 0.49 | 0.13 | 0.69 | 0.16 | 0.73 | 0.07 | 1.51*** | 0.13 |  |  |  |  |
| Ct.Th (mm) | 0.11 | 0.03 | 0.14 | 0.03 | 0.19 | 0.00 | 0.29 | 0.03 |  |  |  |  |
| pMOI (mm<sup>4</sup>) | 0.21 | 0.07 | 0.34 | 0.09 | 0.26 | 0.06 | 0.61** | 0.23 |  |  |  |  |
| TMD (mgHA/cm<sup>3</sup>) | 895 | 44.7 | 878 | 37.3 | 1084 | 5.02 | 99*** | 15.1 |  |  |  |  |
| **Tibia Metaphyseal Bone** |  |  |  |  |  |  |  |  |  |  |  |  |
| TV (mm<sup>3</sup>) | 1.28 | 0.16 | 1.51 | 0.15 | 1.28 | 0.06 | 1.54 | 0.25 |  |  |  |  |
| BV (mm<sup>3</sup>) | 0.46 | 0.15 | 0.65 | 0.21 | 0.50 | 0.06 | 0.83 | 0.19 |  |  |  |  |
| BV/TV (mm<sup>3</sup>/mm<sup>3</sup>) | 0.36 | 0.09 | 0.43 | 0.11 | 0.39 | 0.05 | 0.53 | 0.07 |  |  |  |  |
| vBMD (mgHA/cm<sup>3</sup>) | 246 | 57.7 | 280 | 61.0 | 303 | 42.4 | 404 | 68.4 |  |  |  |  |
| **Femur Metaphyseal Bone** |  |  |  |  |  |  |  |  |  |  |  |  |
| TV (mm<sup>3</sup>) | 1.64 | 0.14 | 2.10*** | 0.18 | 1.59 | 0.07 | 2.23*** | 0.14 |  |  |  |  |
| BV (mm<sup>3</sup>) | 0.59 | 0.20 | 0.83 | 0.32 | 0.59 | 0.05 | 1.14* | 0.18 |  |  |  |  |
| BV/TV (mm<sup>3</sup>/mm<sup>3</sup>) | 0.35 | 0.10 | 0.39 | 0.13 | 0.37 | 0.04 | 0.51 | 0.05 |  |  |  |  |
| vBMD (mgHA/cm<sup>3</sup>) | 246 | 64.1 | 266 | 86.5 | 304 | 26.1 | 385 | 46.3 |  |  |  |  |
|  | 18 wk |  |  |  | 24 wk |  |  |  | 32 wk |  |  |  |
|  | DFF |  | DCKO |  | DFF |  | DCKO |  | DFF |  | DCKO |  |
|  | Mean | St.Dev. | Mean | St.Dev. | Mean | St.Dev. | Mean | St.Dev. | Mean | St.Dev. | Mean | St.Dev. |
| **Tibia Cortical Bone** |  |  |  |  |  |  |  |  |  |  |  |  |
| T.Ar (mm<sup>2</sup>) | 1.17 | 0.17 | 2.39*** | 0.26 | 1.18 | 0.18 | 2.41*** | 0.37 | 1.29 | 0.17 | 2.47*** | 0.22 |
| B.Ar (mm<sup>2</sup>) | 0.74 | 0.10 | 1.8*** | 0.23 | 0.78 | 0.10 | 1.95*** | 0.44 | 0.84 | 0.09 | 2.11*** | 0.48 |
| Ct.Th (mm) | 0.20 | 0.01 | 0.33** | 0.05 | 0.21 | 0.01 | 0.38*** | 0.13 | 0.22 | 0.01 | 0.45*** | 0.10 |
| pMOI (mm<sup>4</sup>) | 0.27 | 0.08 | 1.04*** | 0.21 | 0.28 | 0.09 | 1.17*** | 0.19 | 0.30 | 0.06 | 1.23*** | 0.26 |
| TMD (mgHA/cm<sup>3</sup>) | 1122 | 14.3 | 1025*** | 27.8 | 1133 | 20.9 | 1059*** | 20.2 | 1185 | 19.2 | 1060*** | 20.7 |
| **Tibia Metaphyseal Bone** |  |  |  |  |  |  |  |  |  |  |  |  |
| TV (mm<sup>3</sup>) | 1.33 | 0.10 | 1.80** | 0.28 | 1.39 | 0.24 | 1.86*** | 0.34 | 1.69 | 0.13 | 2.26* | 0.28 |
| BV (mm<sup>3</sup>) | 0.51 | 0.05 | 1.12*** | 0.30 | 0.57 | 0.17 | 1.38*** | 0.44 | 0.67 | 0.13 | 1.88*** | 0.22 |
| BV/TV (mm<sup>3</sup>/mm<sup>3</sup>) | 0.39 | 0.05 | 0.62** | 0.14 | 0.41 | 0.07 | 0.73*** | 0.17 | 0.40 | 0.06 | 0.85*** | 0.17 |
| vBMD (mgHA/cm<sup>3</sup>) | 343 | 57.6 | 539* | 147 | 378 | 70.4 | 689*** | 208 | 417 | 88.8 | 878*** | 181.4 |
| **Femur Metaphyseal Bone** |  |  |  |  |  |  |  |  |  |  |  |  |
| TV (mm<sup>3</sup>) | 1.71 | 0.19 | 2.27*** | 0.22 | 1.74 | 0.18 | 2.31*** | 0.16 | 1.64 | 0.11 | 2.37*** | 0.25 |
| BV (mm<sup>3</sup>) | 0.61 | 0.11 | 1.31*** | 0.33 | 0.64 | 0.14 | 1.53*** | 0.40 | 0.64 | 0.12 | 2.16*** | 0.14 |
| BV/TV (mm<sup>3</sup>/mm<sup>3</sup>) | 0.35 | 0.05 | 0.58** | 0.13 | 0.37 | 0.07 | 0.67*** | 0.17 | 0.39 | 0.05 | 0.92*** | 0.05 |
| vBMD (mgHA/cm<sup>3</sup>) | 312 | 37.6 | 493* | 134 | 338 | 54.5 | 634*** | 204 | 386 | 28.1 | 948*** | 63.4 |

\*p<0.05, \*\*p<0.01, \*\*\*p<0.001 DCKO vs. DFF, Bonferroni's multiple comparisons
#DFF, Fgfr1/2 DFF; DCKO, OC-Fgfr1/2 DCKO

**Supplemental Table S2.** Biomechanics Analysis of femur at 24 wk

|  | DFF |  | DCKO |  |
| --- | --- | --- | --- | --- |
|  | Mean | St.Dev. | Mean | St.Dev. |
| **3-point bending** |  |  |  |  |
| Stiffness | 120 | 41.9 | 175*** | 35.3 |
| Maximum load | 24.1 | 2.32 | 43.8*** | 2.76 |
| post-yield displacement | 0.192 | 0.122 | 0.056* | 0.033 |
| work-to fracture | 6.28 | 1.92 | 7.44 | 2.10 |
| yield load | 21.3 | 2.62 | 40.1*** | 2.52 |
| Young's modulus | 5969 | 1884 | 1797*** | 413 |
| Yield stress | 174 | 29.9 | 98.0*** | 9.92 |
| Maximum stress | 196 | 19.3 | 107*** | 7.35 |
| Moment of inertia | 0.144 | 0.025 | 0.711*** | 0.107 |
| **RPI Testing** |  |  |  |  |
| First unloading slope (US 1<sup>st</sup>) | 0.248 | 0.025 | 0.226 | 0.020 |
| First indentation distance (ID 1<sup>st</sup>) | 37.1 | 7.69 | 40.3 | 5.45 |
| Total indentation distance (TID) | 37.6 | 3.97 | 44.0 | 6.60 |
| Initial indentation distance (IDI) | 6.28 | 0.778 | 8.33 | 2.44 |
| Cumulative indentation distance (CID) | 1.13 | 0.137 | 1.42 | 0.540 |

\*p<0.05, \*\*p<0.01, \*\*\*p<0.001 DCKO vs. DFF unpaired t-test

**Supplemental Table S3.**
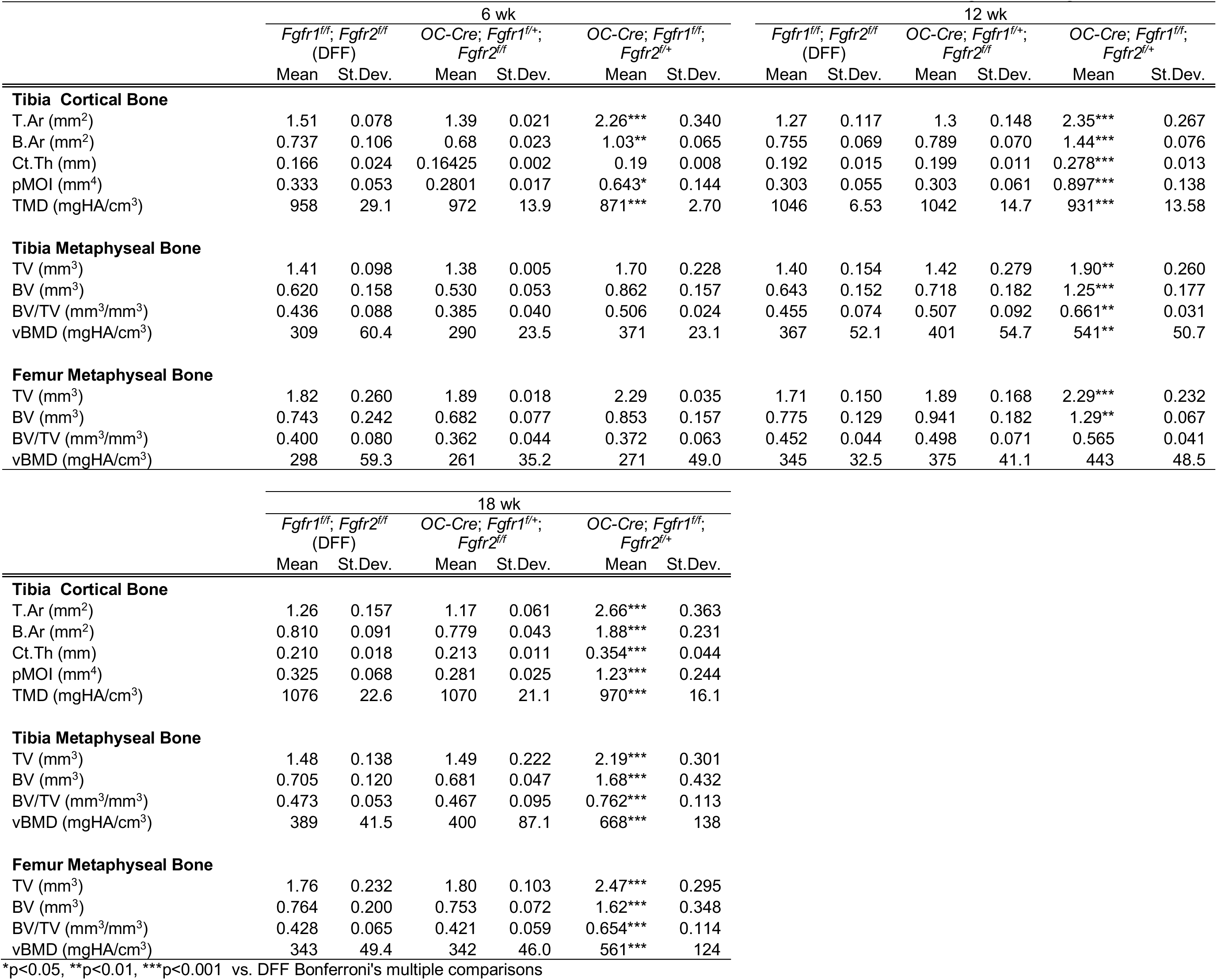
MicroCT analysis of tibia and femur from different allelic combinations of *Fgfr1* and *Fgfr2*

**Supplemental Table S4.** MicroCT analysis of tibia and femur from Fgfr1^f/f^; Fgfr2^f/f^ (DFF) and Dmp1-CreER; Fgfr1^f/f^; Fgfr2^f/f^ (Dmp1-DCKO) mice

|  | 16 wk |  |  |  |
| --- | --- | --- | --- | --- |
|  | *Fgfr1<sup>fl/fl</sup>; Fgfr2<sup>fl/fl</sup>*<br>(DFF) |  | *Dmp1-CreER;*<br>*Fgfr1<sup>fl/fl</sup>; Fgfr2<sup>fl/fl</sup>* |  |
|  | Mean | St.Dev. | Mean | St.Dev. |
| **Tibia Cortical Bone** |  |  |  |  |
| T.Ar (mm<sup>2</sup>) | 1.15 | 0.088 | 1.60** | 0.238 |
| B.Ar (mm<sup>2</sup>) | 0.74 | 0.075 | 1.23* | 0.324 |
| Ct.Th (mm) | 0.198 | 0.021 | 0.263* | 0.048 |
| pMOI (mm<sup>4</sup>) | 0.263 | 0.054 | 0.476* | 0.152 |
| TMD (mgHA/cm<sup>3</sup>) | 1074 | 18.2 | 1032* | 18.0 |
| **Tibia Metaphyseal Bone** |  |  |  |  |
| TV (mm<sup>3</sup>) | 1.73 | 0.203 | 1.99 | 0.249 |
| BV (mm<sup>3</sup>) | 0.682 | 0.055 | 0.819 | 0.194 |
| BV/TV (mm<sup>3</sup>/mm<sup>3</sup>) | 0.398 | 0.042 | 0.041 | 0.048 |
| vBMD (mgHA/cm<sup>3</sup>) | 335 | 53.5 | 335 | 37.2 |
| **Femur Metaphyseal Bone** |  |  |  |  |
| TV (mm<sup>3</sup>) | 1.89 | 0.126 | 1.83 | 0.133 |
| BV (mm<sup>3</sup>) | 0.711 | 0.126 | 0.663 | 0.021 |
| BV/TV (mm<sup>3</sup>/mm<sup>3</sup>) | 0.378 | 0.065 | 0.365 | 0.037 |
| vBMD (mgHA/cm<sup>3</sup>) | 323 | 49.1 | 321 | 34.4 |

\*p<0.05, \*\*p<0.01, \*\*\*p<0.001 vs. DFF unpaired t-test

